# Sticker-and-Linker Model for Amyloid Beta Condensation and Fibrillation

**DOI:** 10.1101/2022.06.04.494837

**Authors:** Jack P. Connor, Steven D. Quinn, Charley Schaefer

## Abstract

A major pathogenic hallmark of Alzheimer’s disease is the presence of neurotoxic plaques composed of amyloid beta (A*β*) peptides in patients’ brains. The pathway of plaque formation remains elusive, though some clues appear to lie in the dominant presence of A*β*_1–42_ in these plaques despite A*β*_1–4_ making up approximately 90% of the A*β* pool. We hypothesise that this asymmetry is driven by the hydrophobicity of the two extra amino acids that are incorporated in A*β*_1–42_. To investigate this hypothesis at the level of single molecules, we have developed a molecular ‘sticker-and-linker lattice model’ of unfolded A*β*. The model protein has a single sticker that may reversibly dimerise and elongate into semi-flexible linear oligomers. The growth is hampered by excluded-volume interactions that are encoded by the hydrophilic linkers but is rendered cooperative by the attractive interactions of hydrophobic linkers. For sufficiently strong hydrophobicity, the chains undergo liquid-liquid phase-separation (LLPS) into condensates that facilitate the nucleation of fibres. We find that a small fraction of A*β*_1–40_ in a mixture of A*β*_1–40_ and A*β*_1–42_ shifts the critical concentration for LLPS to lower values. This study provides theoretical support for the hypothesis that LLPS condensates act as a precursors for aggregation and provides an explanation for the A*β*_1–42_-enrichment of aggregates in terms of hydrophobic interactions.

## I. INTRODUCTION

Alzheimer’s disease (AD), the most common cause of dementia (60%-80% of cases [1]), is a fatal neurodegenerative disease causing sever and devastating cognitive impairment. Age is the biggest risk factor for AD, of which, most cases are sporadic (around 95%) and occur over the age of 65 [1–4]. The exact cause of AD is not fully understood and with better living conditions meaning average life expectancy in developed countries is increasing, the number of cases and the burden of neurodegenerative disease is increasing worldwide. As of 2018, an estimated 50 million people worldwide suffer from dementia, with this number expected to triple by 2050 [5].

A major pathogenic hallmark in AD brains is the presence of extracellular neurotoxic plaques made up of amyloid beta (Aβ) [6, 7]. A*β* is produced from amyloid precursor protein (APP) which is cleaved by *α*, *β* and *γ* secretases [8–10]. APP proteolytic cleavage can be separated into non-amyloidogenic and amyloidogenic pathways. We will focus on the amyloidogenic pathway as it is relevant to AD. First APP is cleaved by *β* secretase into soluble APP*β* and a 99 amino acid C-terminal fragment (C99). C99 is then cleaved by *γ* secretase at multiple sites giving rise to A*β* of multiple lengths ranging from 39-51 amino acids [11, 12]. After cleavage, A*β* is secreted from the cell where it can then form oligomers, fibrils and finally plaques. The amyloid cascade hypothesis suggests that the deposition of A*β* into senile plaques is critical for AD pathology [13–15] but the exact mechanism and change in AD brains that lead to this neurotoxic aggregation is not fully understood. Recently, this hypothesis has been called into question based on amounting evidence in disagreement. Amyloid deposition does not correlate with neuronal loss [16] and amyloid burden can be identified in cognitively unimpaired individuals [17, 18]. Furthermore, clinical trials using therapeutics that target A*β* for degradation have been ineffective thus far [19]. A particular issue for anti-Aβ therapeutics is that most treatments focus on clearing insoluble aggregates [20] with increasing evidence suggesting that pre-fibrillar soluble oligomers are orders of magnitude more toxic than fibrils and plaques [21–24]. Unlike plaques, oligomers have been shown to impair both synaptic function and structure [16, 25]. Recent work by [26] have demonstrated that transthyretin is neuroprotective which is achieved by binding to A*β* oligomers, inhibiting primary and secondary nucleation without altering elongation, ephasising the role of oligomers in A*β*-mediated neurotoxicity. Consequently, understanding the role of pre-fibrillar A*β* species such as soluble oligomers and their contribution to AD pathology has gained significant interest in recent years.

The most predominant A*β* species is 40 amino acids long (A*β*_1–40_) and makes up 90% of the A*β* pool. A less prevalent 42 amino acid long species of A*β* (A_β1-42_), 10% of the A*β* population, is of particular importance to AD due to a higher aggregation propensity. Compared to A*β*_1–40_, A*β*_1–42_ has a longer chain length that increases a repulsive excluded-volume interaction, but adds an attractive hydrophobic interaction due to the addition of two C-terminal hydrophobic amino acids. As such, A*β* _1-42_ has increased aggregation propensity and is a predominant component of senile plaques [27–29]. Familial AD mutations in APP, PSEN1 and PSEN2, cause an increased production of A*β*, with abnormally high levels of A*β*_1–42_ relative to A*β*_1–40_ [30–33]. Recent work shows that the ratio of A*β*_1–42_:A*β*_1–40_ in plasma and cerebrospinal fluid can be used as a biomarker for AD diagnosis [34–40] as it correlates with A*β* deposition [41] and cognitive decline [42–45]. This evidence suggests that an increased ratio of A*β*_1–42_:A*β*_1–40_ may promote the formation of neurotoxic aggregation and the progression of AD. We hypothesise that when A*β*_1–42_ interacts with A*β*_1–40_ sufficient hydrophobicity is added to enhance self-assembly, while the excluded volume interaction is limited. We will model this on a molecular level by describing the intrinsically disordered protein as chains with hydrophilic and hydrophobic segments, as well as ‘sticky’ building blocks that may reversibly self-ensemble into needle-like aggregates.

The aggregation of intrinsically disordered polypeptides (IDP) is an important topic in the biological physics of vital intra- and extracellular processes. In particular, this class of proteins is known to undergo liquid-liquid phase separation (LLPS) to serve biological functionality, such as the (temporary) formation of membraneless organelles [46, 47], but is also appears responsible to form large droplet-like precursors for the aggregation of the tau protein [48], which is another hallmark for Alzheimer’s disease. Large structures (possibly micellar) of approximately 50 A*β* molecules that are speculated to facilitate A*β* aggregation have also been observed [48]. In the present work, we analyse the self-assembly of single peptides to address the current lack of physical models that explain both the formation of large agglomerates and the transition into fibrillar structures in terms of the molecular properties of the IDPs.

We argue that the general mechanism of coupled LLPS and aggregation is an example of physics that emerges from simple concepts such as (non-specific) hydrophobic interactions, excluded volume, and specific interactions. This is a hopeful scenario, as it implies that we may gain relevant molecular insight into this mechanism using strongly coarse-grained molecular models, rather than full atomistic molecular-dynamics simulations that are computationally too demanding to reach the relevant timescales. A commonly used coarse-graining approach to capture both the polymer physics of the polypeptide and the localised formation of reversible non-covalent bonds is to develop a sticker-and-linker model. Such models where originally developed for synthetic associating polymers [49], and were later adopted for disordered proteins, including natural silk [50, 51] and scaffolding proteins in biological condensates, see e.g., [46, 52]. Typical simulation approaches solve the Brownian dynamics of the proteins using simulation software packages such as MARTINI or LAMMPS. A major challenge, which is receiving wide attention from the modelling community [51, 53-55], remains to couple the (continuum-time) conformational dynamics to model the stochastic (instantaneous) association and dissociation of reversible bonds between the molecules. To circumvent the current methodological challenges, in the present work we will employ the fully stochastic Bond-Fluctuation Model [56, 57], which by modelling the molecular dynamics on a 3D lattice successfully describes the (self-avoiding) random walk statistic of flexible and persistent chains [58–60], LLPS, [61, 62], ring-polymers [63], and cross-linking reactions [64]. Crucially, as the dynamics is fully stochastic and discrete, they can be addressed using previously developed kinetic Monte Carlo (kMC) schemes [65].

### B. Method: Bond-Fluctuation Model

We model the chain conformations by placing the polymer segments on the sites of a discrete lattice, following the so-called Bond-Fluctuation Model (BFM) [56, 57].

In the following, we will present a simple sticker-and-linker model for the dimerisation and (linear) oligomerisation of unfolded A*β*, where the cooperativity of growth is determined both by the chain conformation and by the binding energies for dimerisation/nucleation, *ε_n_*, and elongation, *ε_e_*, and where hydrophobic interactions are modelled using a short-ranged interaction energy *ε*_H_. Subsequently, we will present the bond-fluctuation model using which we simulate the aggregation of the A*β*. In our analysis, we first investigate how the hydrophobic interactions affect the partial collapse of the molecule, and how it may lead to LLPS. We then show that these interactions facilitate dimerisation, as well as the further cooperative growth into longer oligomers. Lastly, we demonstrate that increasing A*β*_1–42_ relative to A*β*_1–40_ promotes aggregation.

## II. THEORY AND METHOD

### A. Introduction: Parametrisation of unfolded A*β*

A*β* may be viewed as a (partially) intrinsically disordered peptide, given the findings of random coil conformations through both circular dichroism [66] and nuclear magnetic resonance spectroscopy [67, 68]. Hence, we chose to use a model of unfolded monomeric A*β* in agreement with recent biophysical analysis that suggests that A*β* monomers do not adopt a stable secondary structure *in vitro* conditions. We parametrise this model using the sequence of amino acids in A*β*_1–42_, see **Figure 1A**, which displayes the hydrophilic residues in red, the neutral ones in black and the hydrophobic ones in blue. Our model for A*β* (**Figure 1B**) was based on the structure of A*β*_1–42_ in [69]. The first eight grey (inert) beads represent the first 16 amino acids corresponding to the hydrophilic N-terminal region. The central hydrophobic core is modelled as two blue (hydrophobic) beads representing amino acids 17-21. A red ‘sticky’ bead which is capable of dimerising with other red beads was added into the turn region to recapitulate the intra- and interchain salt bridges that can occur in this region. The remaining hydrophobic beads represent the C-terminal region with A*β*_1–42_ containing one extra bead to account for the addition of two hydrophobic amino acids.

**FIG. 1:**
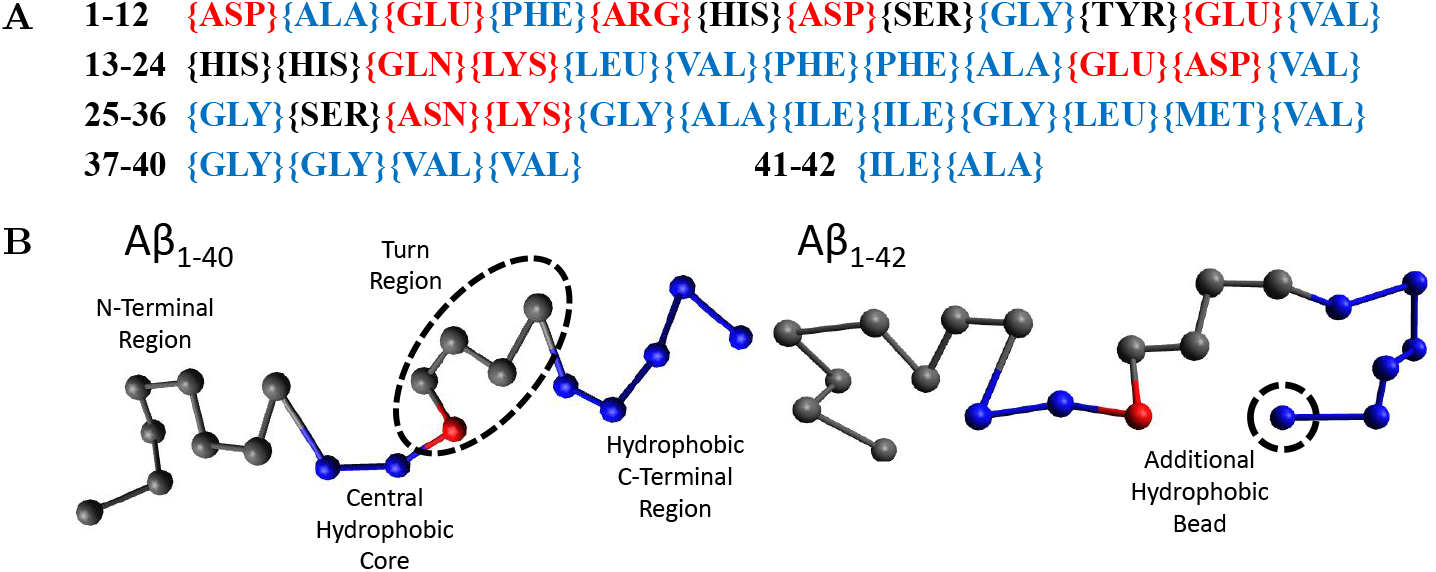
**(A)** The amino acid sequence of A*β*_1–42_. Amino acids labelled red are hydrophilic, black are neutral and blue are hydrophobic. **(B)** Our beads on a string model for both A*β*_1–40_ and A*β*_1–42_ with the N-terminus to the C-terminus going from left to right. Each bead represents 2 amino acids. Grey beads are inert, representing the N-terminus (1-16) and the turn region (24-27). Blue beads are hydrophobic, representing the central hydrophobic core (17-21), the second hydrophobic region (29-35) and the hydrophobic C-terminus (36-40/42). Red beads are capable of dimerising with other red beads.

This approach originates from the success of lattice models to predict phenomena such as LLPS, including in conditions near the critical point where the correlation-length diverges and mean-field models break down [70–72]. Similarly, the lattice models have provided computationally efficient means to sample the configuration space of polymers. Early lattice models for polymers place polymer segments on a single lattice site [73], but underestimated the Rouse dynamics of the chains due to kinetically trapped states. This was was remedied in the BFM by letting a segment occupy 8 sites on a 3D cubic lattice (4 sites on a 2D square lattice) and where the bond lengths can fluctuate from 2 to 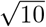 in lattice units [56, 57]. This approach was proven competitive with off-lattice models both in terms of computational convenience and in physical accuracy in (and beyond) the examples mentioned in the Introduction [58, 61–64].

Within the BFM, the polymer conformations and/or dynamics are modelled using kinetic Monte Carlo (kMC) time steps in which a monomer may move in 6 directions on the lattice if this does not (1) overstretch or understretch the bond between two monomers within the chain or (2) lead to double-occupied lattice sites. This leads to a list of *N*_enabled_ ≤ 6*N*_monomers_ possible processes that may occur during the time step, of which only one (or none) may take place. Which process, *i,* may take place is selected using the rate,

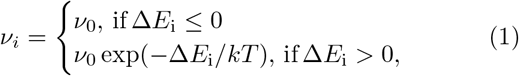

where *ν*_0_ is an elementary rate (typically of the order 1 – 100*μ*s^−1^) and where Δ*E*_i_ is the change in energy. In the algorithm that we use, a process is randomly selected out of the *N*_enabled_ list of processes. The process is then executed if the system energy is unchanged or decreases Δ*E*_i_ ≤ 0. However, if the energy would increase, the process is only executed with a probability *p* = exp(–Δ*E*_i_/*kT*) and rejected otherwise; this decision is carried out using random numbers drawn using a SIMD-oriented Fast Mersenne Twister[74]. Regardless if the process is accepted or rejected, the time is increased with Δ*t* = 1/(*N*_enabled_*ν*_0_). The computational efficiency of the method relies on the fact that following a Monte Carlo step during which a monomer moves, only the rates of monomers in the vicinity of this monomer needs to be recalculated [65].

### C. Parametrisation: Hydrophobicity of Linkers

To model the attraction between two hydrophobic monomers we follow the approach by Reister et al., and use a square-well potential [61]

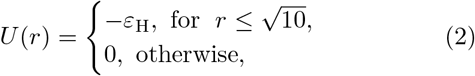

where *r* is the distance (in units of the lattice spacing) between the two monomers, and where ε_H_ describes nonspecific (e.g., hydrophobic) interactions [61]. This interaction energy enables the parametrisation of intrinsically disordered poly-peptides through the radius of gyration, and may in principle depend on conditions such the temperature and the ionic strength [75, 76]. This parametrisation is done using the Flory exponent ν, which describes the swelling of a polymer through the radius of gyration as 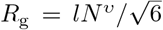, with *l* the step length between segments. In good solvent conditions, the chain is swollen due to intramolecular self-excluded volume interactions and *ν* = 0.588. Completely insoluble chains, described with a large value of *ε*_H_, collapse to a compact sphere with *ν* = 1/3. At *θ* conditions the hydrophobic interactions exactly cancel the excluded-volume interactions and the chain obeys random-walk statistics, which are characterised by *ν* = 1/2. To find the *θ*-condition, we measure the radius of gyration for chains with various chain length, *N*, as a function of *ε*_H_ [77, 78]. As 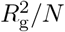 is independent of the chain length at the theta condition, the *ε*_H_ value at which all curves intersect represents the *θ* condition. From **Figure 2**, we find that this occurs at *ε*_H_/*k*_B_*T* ≈ 0.27 for (homopolymer) chains with identical subunits.

**FIG. 2:**
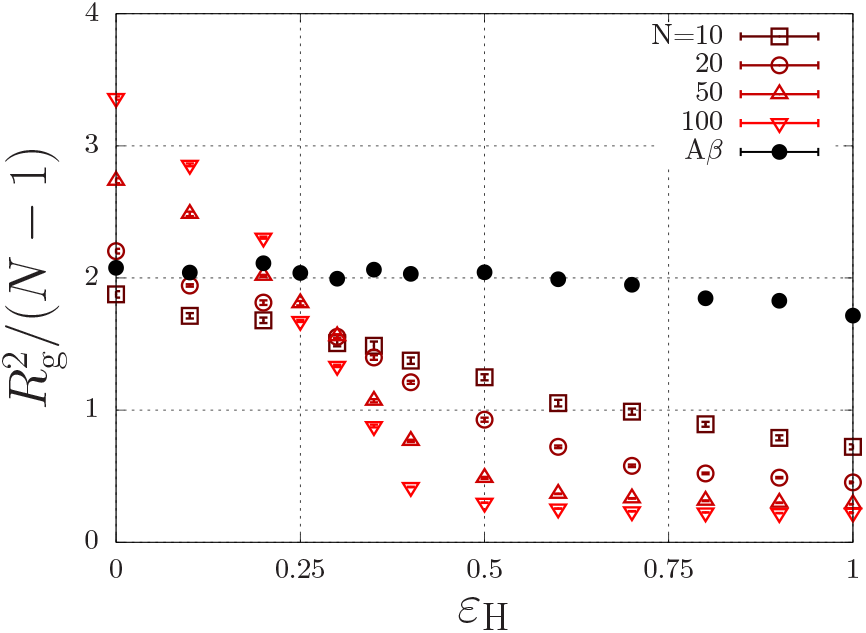
Polymer size, 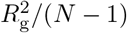, (*R*_g_ is the radius of gyration in units of the lattice spacing) as a function of the hydrophobic interaction energy *ε*_H_/*k*_B_*T* for a various number of beads per chain, *N*. The A*β* model has of total of *N* = 20 beads, of which 7 are hydrophobic.

As discussed in the previous section, our model for A*β* describes the protein as a copolymer with both hydrophilic and hydrophobic units. The solid circles in **Figure 2** show that the radius of gyration, 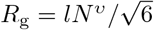 of this model polypeptide is relatively insensitive of the hydrophobic interaction parameter. To estimate the overlap concentration above which the intramolecular excluded-volume interactions are screened, we use the end-to-end distance, given by *R*_e_ = *lN^v^* ≈ 3.3 nm, where A*β* has *N* = 40 amino acids, and where *l* = 0.36 nm is the typical step length of amino acids [75, 76]. Using the molecular volume 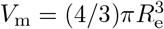, we find an overlap concentration of *c*\* = *M*_w_/(*N*_A_*V*_m_) ≈ 100 mg/ml = 0.025 M, where we used *M*_w_ = 4.5 kg/mol. In typical experiments, aggregation is observed well below the overlap concentration (e.g., A*β*_1–40_ aggregates below 1 mg/ml [79] and A*β*_1–42_ aggregates at concentrations as low as 90 nM [80]).

While those experimental conditions may suggest dilute conditions in which excluded-volume interactions are unimportant, inside LLPS condensates the concentration is signicantly higher, and may in fact exceed the overlap concentration. To identify LLPS in our simulations, we focus on structural coarsening phenomena such as Ostwald or Lifshitz-Slyozov-Wagner (LSW) ripening and/or Brownian coalescence [81]. The dynamics of coarsening is typically associated with a characteristic length scale (e.g., the radius of a droplet) that grows with the one-third power of time as *R*\* – *R*^0^ ∝ *t*^1/3^ with *R*_0_ the initial length scale, which may emerge through nucleation or spinodal decomposition. In our simulations, we determine *R*\* using the following recipe [71, 72, 82]: For a given time, *t*, we calculate the order-parameter field *ψ*(**r**_*i*_), with **r** the spatial coordinate of a lattice site with i the index of a lattice site. The value of *ψ*(**r**_*i*_) is set to 1 if the site is occupied by a hydrophobic monomer, and to –1 otherwise. We then calculate the 3D Fourier transform 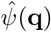, and obtain the structure factor 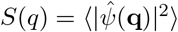, where 〈·〉 is the spherical/angular average. Next, we obtain the spatial correlation function *C*(*R*) as the inverse Fourier transform of *S*(*q*). As discussed previously[71, 72, 82], numerically robust measures for the characteristic length scale *R*\* are the first root (i.e, given by *C*(*R*\*) = 0) and the first minimum for which *C*(*R*\*) < 0 and *C*′(*R*\*) = 0. In our analysis we will use both measures to assess if structural coarsening occurs in our simulations.

### D. Parametrisation: Aggregation of Stickers

Assuming that the formation of A*β* aggregates is, in principle, a reversible process, we consider two aggregates of size *m* ≥ 1 and *n* ≥ 1 that may reversibly assemble into an aggregate of size *m* + *n* through the chemical equilibrium equation [83–85]

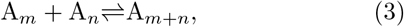

independent of the molecular configuration; i.e., an aggregate larger than *n* > 2 may be a linear oligomer or a ring [86], or a species with a different topology. In our modelling, we define binding energies that can only lead to dimers, linear oligomers, and rings. To focus on the linear oligomerisation of A*β*, rather than ring formation, we will also define bending potentials that are strong enough to prevent ring formation but are sufficiently weak to avoid lattice-imposed alignment artifacts.

As dimerisation often appears to act as a nucleation step in the process of oligomerisation [85, 87], we define different binding energies for the dimerisation and the elongation steps. The dimerisation reaction is

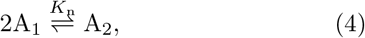

where

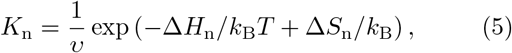

is the equilibrium constant for nucleation, with *ν* some characteristic volume, and Δ*H* < 0 is the binding enthalpy and with Δ*S*_n_ > 0 the entropic penalty of dimerisation. In this equation, Δ*H*_n_ is controlled by the binding energy *ε*_n_ > 0 and the hydrophobic interaction strength *ε*_H_. In the absence of hydrophobic interactions, Δ*H*_n_ = –*ε*_n_ is exact. The entropic penalty essentially originates from an excluded-volume interaction due to a limitation of the internal degrees of freedom (DOF) of two chains that undergo dimerisation, see section III A. In that section we will show that these internal DOF are affected by the hydrophobicity (an increased hydrophobicity partially collapses the chain and limits the DOF prior to dimerisation), and by the concentration (above the overlap concentration the free chains have fewer DOF prior to dimerisation), such that an increasing hydrophobicity and concentration reduce the entropic penalty and enhance dimerisation.

For subsequent oligomerisation steps, i.e., for *n* > 1, we describe the equilibrium statistics using

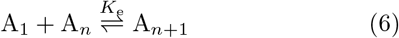

with

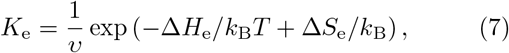

in which Δ*H*_e_/*k*_B_*T* = - *ε*_n_ + *U*_hydrophobic_ + *U_θ_* is composed of an oligomeration/elongation energy, *ε*_n_ > 0, and hydrophobic interactions as before, but is also affected by the conformation of the oligomer: It is fully flexible for angles smaller or equal than *θ*_max_, but infinite for angles larger than that (for applications of smoother potentials in BFM simulations, see [58, 77, 88]).

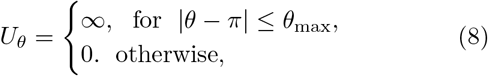

One of the key predictions of our simulations will be the dependence of the fraction of aggregated material on the concentration and hydrophobicity. We will interpret these findings using analytical predictions in the limit where hydrophobic interactions are absent and where the entropic penalties are constant. In this limit there exist some known analytical predictions [84]. The starting point to obtain these is by writing the equilibrium constants as *K*_n_ = [*A*_2_]/[*A*_1_]^2^ and as *K*_e_ = [*A*_*n*+1_]/[*A_n_*][*A*_1_], which are constant for all *n* ≥ 2, so that the concentration of any aggregate [*A*_n_] with *n* ≥ 2, can be expressed in terms of the concentration of unbound A*β*, [A_1_], as

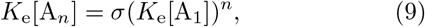

with *σ* ≡ *K*_n_/*K*_e_ the so-called cooperativity factor.

The concentration of unbound A*β* is obtained from the mass balance,

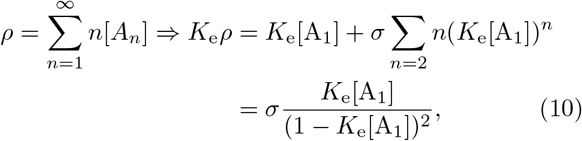

where *ρ* is the (experimentally-controlled) overall concentration of A*β*. Using the standard sum 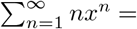

We have curve fitted each data set with a fixed dimerisation energy, *ε*_e_, using the dimerisation model of **Equation (12)** (solid curves) with the equilibrium constant *x*/(1 – *x*)^2^ for |*x*| < 1 and the fraction of aggregated molecules, *f* ≡ 1 – [*A*_1_]/*ρ*, this mass balance can be written as [83, 85]

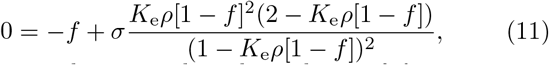

which provides an implicit dependence of *f* on *ρ*.

This equation has three asymptotic limits of interest, namely the strongly anti-cooperative case where dimers do not grow into larger aggregates, *K*_e_ → 0 (resulting in *σ* → ∞) leads to [83]

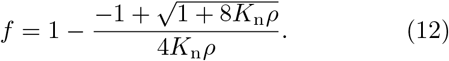

In the non-cooperative case, or ‘isodesmic’ case [89], *K* ≡ *K*_e_ = *K*_n_ [83],

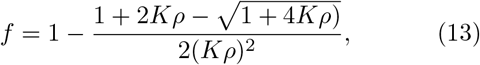

and in the strongly cooperative case, *σ* → 0 [84, 89], we have *f* = 0 for *K*_e_*ρ* < 0 and

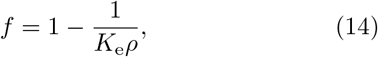

for *K*_e_*ρ* ≥ 0. These equations show that from the anticooperative to the cooperative case, the transition from unbound to aggregated material becomes increasingly sharp. In the following, we will discuss the influence of hydrophobic and excluded-volume interactions on self-assembly in terms of the elongation constant *K*_e_ and the cooperativity factor, *σ*.

## III. RESULTS AND DISCUSSION

### A. Dimerisation and LLPS

To investigate how hydrophobic and steric interactions affect dimerisation, we have compared the self-assembly of chains with such interactions to the completely hydrophylic counterpart with *ε*_H_ = 0, and disabled oligomerisation (i.e., *ε*_e_ = 0). We have then simulated the molecular self-assembly of 100 chains in a periodic simulation box with sizes ranging from 50 × 50 × 50 to 1000 × 1000 × 1000, with dimerisation energies ranging from *ε*_n_ = 2 to 10*k*_B_*T*. The resulting fraction of dimerised chains, *f*, against the number density *ρ* (in number molecules per box size), is represented by the symbols in **Figure 3A**.

**FIG. 3:**
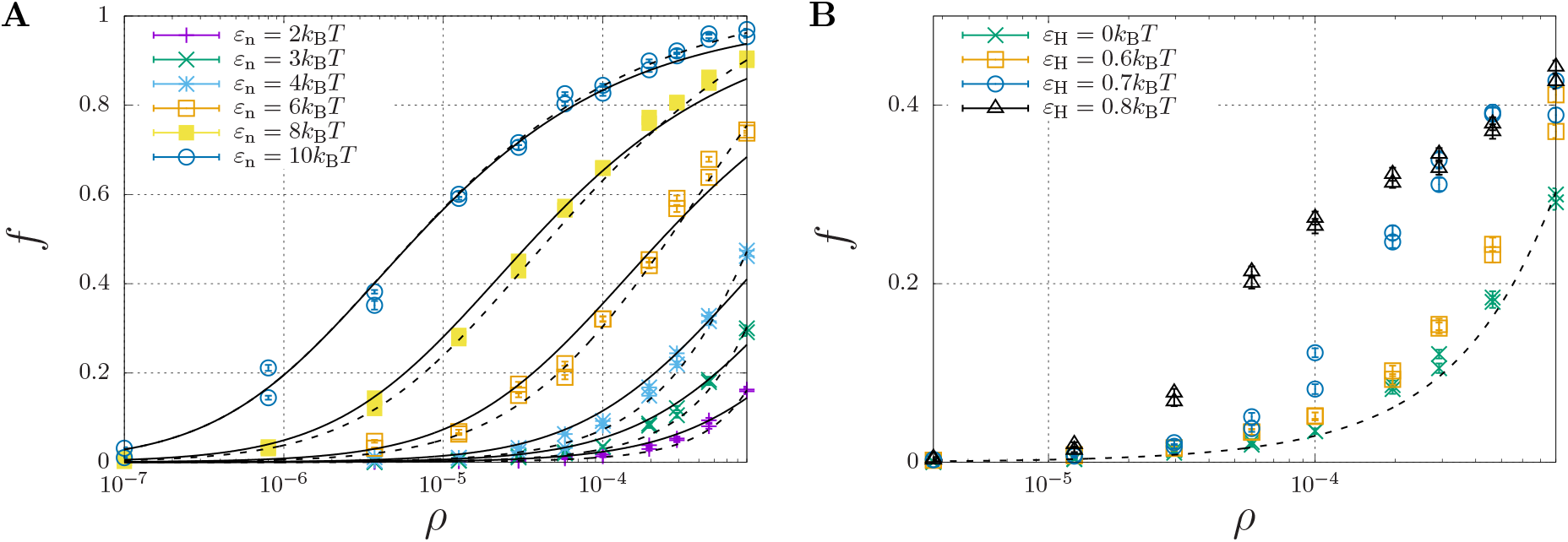
Fraction of dimerised A*β, f*, plotted against the number density, *ρ* (in units of the lattice spacing). Well below the overlap concentration of *ρ* = 10^-4^ (corresponds roughly to 100 mg/ml, see main text) the solution can be considered dilute, while at higher concentrations excluded volume interaction become important. **(A)** The dimerisation energy *ε*_n_ is varied from 2 to 10*k*_B_*T* without hydrophobic interactions (*ε*_H_ = 0). The solid curves are individual fits using **Equation (12)** with a dimerisation constant *K*_n_ = *K*_n,0_ exp(*ε*_n_) with a non-constant *K*_n,0_ (**Table I**), while the dashed curves represent a simultaneous fit of all data using *K*_n,0_ = *K*_n,00_ exp[(*∂*[Δ*S*_n_/*k*_B_]/*∂ρ*)*ρ*] with constant *K*_00_ = 6.85 and an entropic excluded-volume correction (*∂*[Δ*S*_n_/*k*_B_]/*∂ρ*) = 1.3 · 10^3^. **(B)** The dimerisation energy is fixed *ε*_n_ = 3*k*_B_*T*, while the hydrophobicity is increased from 0 up to *ε*_H_ = 0.8. The dashed and dotted curve are calculated using *K*_n_ = 6.85 exp(3 + 1300*ρ*) and *K*_n_ = 6.85 exp(3.6 + 1300*ρ*), respectively. The data for *ε*_H_ = 0.7 and 0.8 are underestimates, as ongoing LLPS enables the slow increase of aggregate sizes.

*K*_n_ = *K*_n,0_exp(*ε*_n_). While *K*_n,0_ is explicitly independent of *ε*_n_, the curve fits yielded the apparent dependence ln *K*_n,0_ ≈ 3.0 – 1.1*ε*_n_ (**Table I**). This is caused by the fact that the dimerisation concentration (characterised by the inflection point of the self-assembly curve) shifts to higher concentrations (above the overlap concentration of *ρ* ≈ 10^-4^, see section II C) for decreasing interaction strengths. Following Flory’s approach to self-avoiding walk statistics, we modify the entropy for excluded-volume interactions using a meanfield description, Δ*S*_n_ = Δ*S*_n,0_ + (*∂*Δ*S*_n_/*∂ρ*)*ρ*, and write *K*_n_ = *K*_n,00_ exp(*ε*_n_ + (*∂*[Δ*S*_n_/*k*_B_]/*∂ρ*)*ρ*. Using this modification, we have been able to capture all dimerisation curves in **Figure 3A** with fixed *K*_00_ = 6.85 and excluded-volume correction (*∂*[Δ*S*_n_/*k*_B_]/*∂ρ*) = 1.3 · 10^3^ (dashed curves). As expected, this correction is insignificant below the overlap concentration, *ρ* < 10^-4^, but significantly affects the self-assembly curves at higher concentrations.

**TABLE I:**
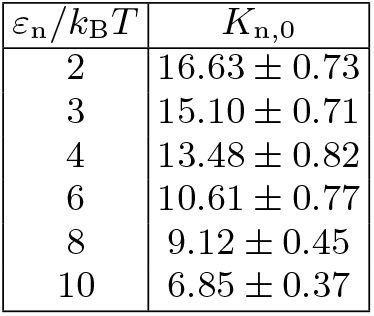
Values for the equilibrium constant *K*_n,0_ determined by curve-fitting **Equation 12** to the simulated data for various values of the dimerisation energy εn. The curve-fits are shown as solid lines in **Figure 3**. A lin-log regression yields an *apparent* relationship ln *K*_n,0_ ≈ 3.0 – 1.1*ε*_n_/*k*_B_*T* (see main text).

We now focus our attention on the dimerisation curve with *ε*_n_ = 3*k*_B_*T*, which has a very low fraction of dimers below the overlap concentration but a finite fraction of dimers at higher concentrations, and increase the hydrophobic-interaction parameter, *ε*_H_, from 0 to 0.8*k*_B_*T* (**Figure 3B**). We find that up to a value 0.6*k*_B_*T*, the hydrophobicity appears to only modestly modify the equilibrium constant as *K*_n_ ≈ 6.85exp(*ε*_n_ + *ε*_H_ + 1300*ρ*); we speculate that only the central hydrophobic beads contribute to the enhancement of dimerisation. For stronger hydrophobic interactions, however, the shape of the self-assembly curve can no longer be described using the simple dimerisation model.

In fact, the datapoint for *ε*_H_ ≥ 0.7 in **Figure 3B** is not fully converged, and the fraction of dimers increases in a slow process, as indicated in **Figure 4A**. This figure shows that the fraction of dimers, *f*, reaches a plateau at *f* ≈ 0.2 for times *t* > 100 up to *t* ≈ 3000, but then slowly increases up to *f* ≈ 0.4 at *t* = 10^5^ without any sign of convergence to a higher plateau value. This process is a consequence of the slow formation of large structures, as indicated by the typical length scales in the system **Figure 4B**, which increase as *R*\* – *R*^0^ ∝ *t*^1/3^, which is a hallmark of LLPS, see section II C. Here, we determined R* using the structure factor *S*(*q,t*) (right top), and its corresponding spatial correlation function, *C*(*R,t*), (right bottom). We quantified *R*\* using the first root (*C*(*R*\*, *t*) =0, *R*^0^ = 8) and the first minimum (*C*(*R*\*, *t*) < 0 and *∂C/∂R* = 0, *R*^0^ = 12).

**FIG. 4:**
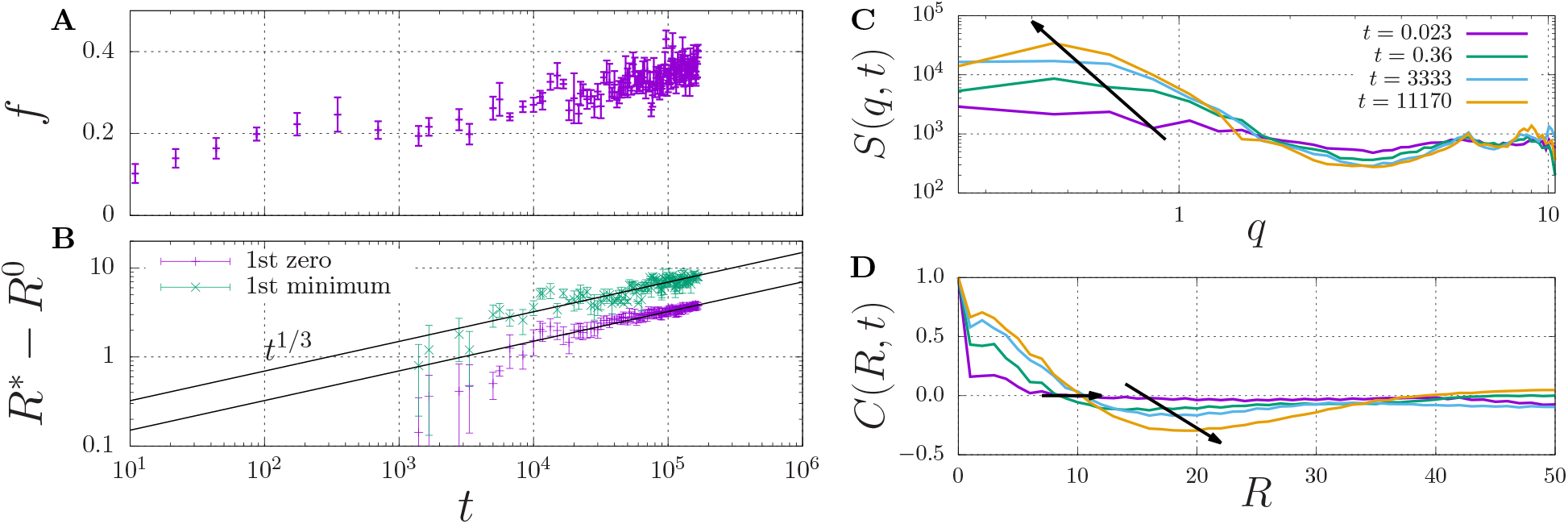
Fraction of dimerised material, *f*, **(A)** and characteristic length scale, *R*\* – *R*^0^ **(B)** against time. The characteristic length scale, *R*\* is determined by i) calculating the dynamic structure factor, *S*(*q*) **(C)**, ii) taking the inverse Fourier transform to obtain the radial correlation function, *C*(*R, t*) **(D)**, and iii) determining its first root *C* = 0 (here we use the initial offset *R*^0^ = 8) and first minimum for which *C* < 0 (here, *R*^0^ = 12). The arrows indicate the time dependence of the dominant wavenumber q **(C)** and the two measures for *R*\* **(D)**.

These findings indicate that sufficiently strong hydrophobic interactions may lead to to formation of droplets through LLPS, which inside the droplets increases the concentration of dimerising units beyond the overlap concentration (which screens the excluded-volume interactions), and provides the mass action needed to induce dimerisation.

### B. Oligomerisation

Now that we have investigated the dimerisation of A*β*, we will investigate the growth into larger oligomers. To that purpose we again first focus on the case without any hydrophobic interactions, as this enables us to isolate the impact of excluded-volume interactions on the (anti-)cooperativity of self-assembly. This also enables us to select nucleation and elongation energies of interest in the subsequent simulations, in which we do switch on hydrophobic interactions. Akin to the dimerisation case, we have simulated the self-assembly of 100 A*β*_1–40_ chains in box sizes ranging from 36 × 36 × 36 to 500 × 500 × 500. In these simulations, we have swithed off the non-specific/hydrophobic interactions *ε*_H_ = 0, and varied the nucleation energies in the range *ε*_n_ = 1 – 9*k*_B_*T* and the elongation energies in the range *ε*_e_ = 8 – 14*k*_B_*T*.

The simulations yielded the fraction of aggregated material, *f*, as a function of the concentration, to which we have curve fitted the theoretical model of **Equation (11)**. From the curve-fits we have extracted the cooperativity factor *σ* and the equilibrium constant *K*_e,0_ as a function of the dimerisation energy. Here, *K*_e,0_ is defined by

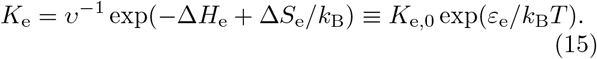

Th results are tabulated in **Table II**) and plotted in **Figure 5B** and **C** (discussed below).

**TABLE II:** Shown are the results from varying *ε*_n_ for multiple different values of *ε*_e_, in the absence of hydrophobic interactions i.e. *ε*_H_ = 0*k*_B_*T*. Each simulation used 100 A*β*_1–40_ chains in box sizes ranging from 36 x 36 x 36 to 500 x 500 x 500. Each value of *σ* and *K*_n,0_ represents the mean and standard error from 3 separate simulations

| $\varepsilon_n/k_B T$ | $\varepsilon_e/k_B T$ | $\sigma$ | $K_{n,0}$ |
| --- | --- | --- | --- |
| 1 | 8 | $2.15 \times 10^{-3} \pm 2.31 \times 10^{-3}$ | $2.24 \times 10^{-4} \pm 1.08 \times 10^{-5}$ |
| 3 | 8 | $5.41 \times 10^{-2} \pm 1.24 \times 10^{-2}$ | $1.82 \times 10^{-4} \pm 8.96 \times 10^{-6}$ |
| 5 | 8 | $3.91 \times 10^{-1} \pm 3.73 \times 10^{-2}$ | $1.26 \times 10^{-4} \pm 5.83 \times 10^{-6}$ |
| 7 | 8 | $1.90 \pm 1.65 \times 10^{-1}$ | $7.29 \times 10^{-5} \pm 4.43 \times 10^{-6}$ |
| 9 | 8 | $8.15 \pm 2.10$ | $3.87 \times 10^{-5} \pm 8.80 \times 10^{-6}$ |
| 1 | 9 | $1.33 \times 10^{-4} \pm 4.68 \times 10^{-4}$ | $3.87 \times 10^{-4} \pm 1.89 \times 10^{-5}$ |
| 3 | 9 | $2.10 \times 10^{-2} \pm 9.29 \times 10^{-3}$ | $3.25 \times 10^{-4} \pm 1.72 \times 10^{-5}$ |
| 5 | 9 | $2.34 \times 10^{-1} \pm 3.38 \times 10^{-2}$ | $2.38 \times 10^{-4} \pm 1.31 \times 10^{-5}$ |
| 7 | 9 | $1.50 \pm 1.54 \times 10^{-1}$ | $1.61 \times 10^{-4} \pm 1.11 \times 10^{-5}$ |
| 9 | 9 | $5.55 \pm 1.54$ | $7.92 \times 10^{-5} \pm 1.86 \times 10^{-5}$ |
| 1 | 10 | $1.00 \times 10^{-5} \pm 1.54 \times 10^{-4}$ | $6.89 \times 10^{-4} \pm 3.19 \times 10^{-5}$ |
| 3 | 10 | $5.34 \times 10^{-3} \pm 3.74 \times 10^{-3}$ | $5.83 \times 10^{-4} \pm 2.60 \times 10^{-5}$ |
| 5 | 10 | $1.53 \times 10^{-1} \pm 3.04 \times 10^{-2}$ | $4.49 \times 10^{-4} \pm 2.87 \times 10^{-5}$ |
| 7 | 10 | $1.08 \pm 1.36 \times 10^{-1}$ | $3.26 \times 10^{-4} \pm 2.55 \times 10^{-5}$ |
| 9 | 10 | $6.02 \pm 1.43$ | $2.10 \times 10^{-4} \pm 4.26 \times 10^{-5}$ |
| 1 | 11 | $1.00 \times 10^{-5} \pm 7.55 \times 10^{-5}$ | $1.15 \times 10^{-3} \pm 3.72 \times 10^{-5}$ |
| 3 | 11 | $9.06 \times 10^{-3} \pm 5.57 \times 10^{-3}$ | $1.05 \times 10^{-3} \pm 5.99 \times 10^{-5}$ |
| 5 | 11 | $8.19 \times 10^{-2} \pm 2.17 \times 10^{-2}$ | $7.98 \times 10^{-4} \pm 5.03 \times 10^{-5}$ |
| 7 | 11 | $8.85 \times 10^{-1} \pm 1.15 \times 10^{-1}$ | $6.62 \times 10^{-4} \pm 5.07 \times 10^{-5}$ |
| 9 | 11 | $4.60 \pm 1.18$ | $4.64 \times 10^{-4} \pm 9.82 \times 10^{-5}$ |
| 1 | 12 | $4.58 \times 10^{-5} \pm 1.79 \times 10^{-4}$ | $2.23 \times 10^{-3} \pm 7.59 \times 10^{-5}$ |
| 3 | 12 | $3.77 \times 10^{-3} \pm 3.18 \times 10^{-3}$ | $1.83 \times 10^{-3} \pm 8.20 \times 10^{-5}$ |
| 5 | 12 | $8.19 \times 10^{-2} \pm 1.61 \times 10^{-2}$ | $1.51 \times 10^{-3} \pm 6.71 \times 10^{-5}$ |
| 7 | 12 | $5.94 \times 10^{-1} \pm 9.53 \times 10^{-2}$ | $1.19 \times 10^{-3} \pm 9.93 \times 10^{-5}$ |
| 9 | 12 | $2.88 \pm 5.63 \times 10^{-1}$ | $7.77 \times 10^{-4} \pm 1.15 \times 10^{-4}$ |
| 1 | 14 | $5.95 \times 10^{-3} \pm 5.23 \times 10^{-3}$ | $8.12 \times 10^{-3} \pm 5.12 \times 10^{-4}$ |
| 3 | 14 | $1.65 \times 10^{-2} \pm 1.14 \times 10^{-2}$ | $7.99 \times 10^{-3} \pm 6.79 \times 10^{-4}$ |
| 5 | 14 | $7.83 \times 10^{-2} \pm 2.53 \times 10^{-2}$ | $4.97 \times 10^{-3} \pm 4.04 \times 10^{-4}$ |
| 7 | 14 | $3.26 \times 10^{-1} \pm 6.66 \times 10^{-2}$ | $3.43 \times 10^{-3} \pm 3.03 \times 10^{-4}$ |
| 9 | 14 | $7.21 \times 10^{-1} \pm 1.36 \times 10^{-1}$ | $2.00 \times 10^{-3} \pm 2.07 \times 10^{-4}$ |

**FIG. 5:**
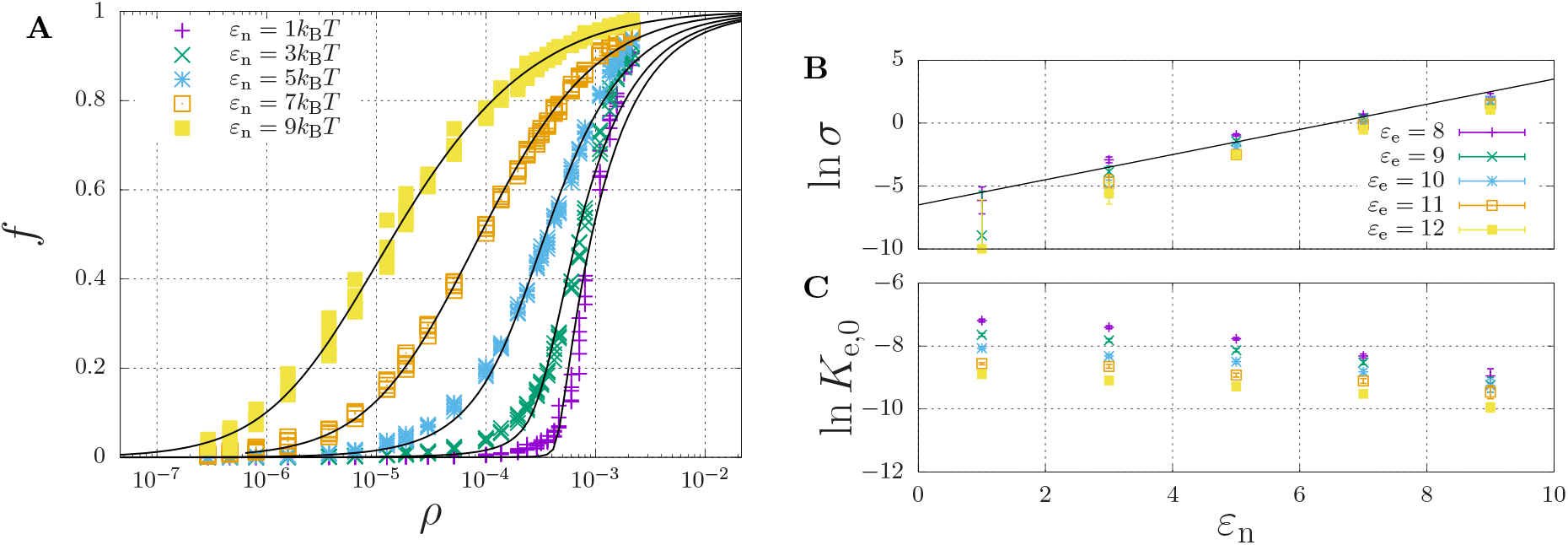
Self-assembly of a hydrophilic chain (*ε*_H_ = 0) for a fixed elongation energy *ε*_e_ = 9*k*_B_*T* and a varying dimerisation energy *ε*_n_. **(A)** The fraction of aggregated material plotted against the concentration. For each value of *ε*_n_ the data was curve-fitted using **Equation (11)** with cooperativity factor *σ* **(B)** and the equilibrium constant *K*_e_ = *K*_e,0_ exp(*ε*_e_) **(C)** as fitting parameters. For **(B)** and **(C)** each point represents the mean and standard error of 5 simulations.

A representative self-assembly curve, obtained for fixed *ε*_e_ = 9*k*_B_*T*, is shown in **Figure 5A**. This panel shows the fraction of aggregated material, *f*, increases with an increasing number density, *ρ*. The sigmoidal curve becomes increasingly sharp with a decreasing dimerisation energy, which, as expected (see section II D), indicates an increasing cooperativity of self-assembly. Indeed, **Figure 5B** shows that the logarithm of the cooperativity factor increases linearly with an increasing dimerisation energy, as expected. On the other hand, the elongation constant *K*_e,0_ is in principle expected to be independent of the dimerisation energy; however, **Figure 5C** shows it apparently decreases with an increasing dimerisation energy. We attribute this apparent dependence to the excluded-volume interactions becoming more present at higher concentrations, as we found above in the dimerisation curves of **Figure 3**.

To investigate the effect of non-specific hydrophobic interaction on oligomerisation, we have used the parameters *ε*_n_ = *3*k_B_*T* and *ε*_e_ = 9*k*_B_*T* of a hydrophylic chain that cooperativily self-assembles at reasonably high concentrations. This ensured that low-concentration structuring due to hydrophobic interactions would not neccessitate larger (and computationally expensive) box sizes. We have then used the sequences for the A*β*_1–40_ and A*β*_1–42_ chains and varied the hydrophobic interaction energy *ε*_H_ from 0.4 to 0.7*k*_B_*T*. We have again simulated 100 chains in box sizes ranging from 36 × 36 × 36 to 500 × 500 × 500 lattice sites.

We have presented the results in **Figure 6**, where panels **A** and **B** display the fraction of aggregated material *f* against the number density *ρ* for A*β*_1–40_ (**A**) and A*β*_1–42_ (**B**). In line with our results on dimerisation, we find that an increasing hydrophobic interaction energy shifts the aggregation concentration to lower concentration, and that the transition to the aggregates state becomes sharp. For these higher hydrophobicities, we again find a slow ongoing increase of the fraction of aggregated material due to Ostwald ripening and/or rare events of fusion of condensates. After close inspection of panels **A** and **B**, we observe that the transition occurs for A*β*_1–42_ at lower concentrations than for A*β*_1–40_, which we attribute to a larger number of hydrophobic interactions for the longer chain. Indeed, **Figure 6C** shows that the mean aggregate size at *ρ* = 2 · 10^-4^ sharply increases at *ε*_H_ = 0.6 for the longer chain and at *ε*_H_ = 0.65 for the shorter chain. The mean values of the aggregate size are biased by the presence of small oligomers; **Figure 6D** reveals the presence of aggregates with 8 or more chains.

**FIG. 6:**
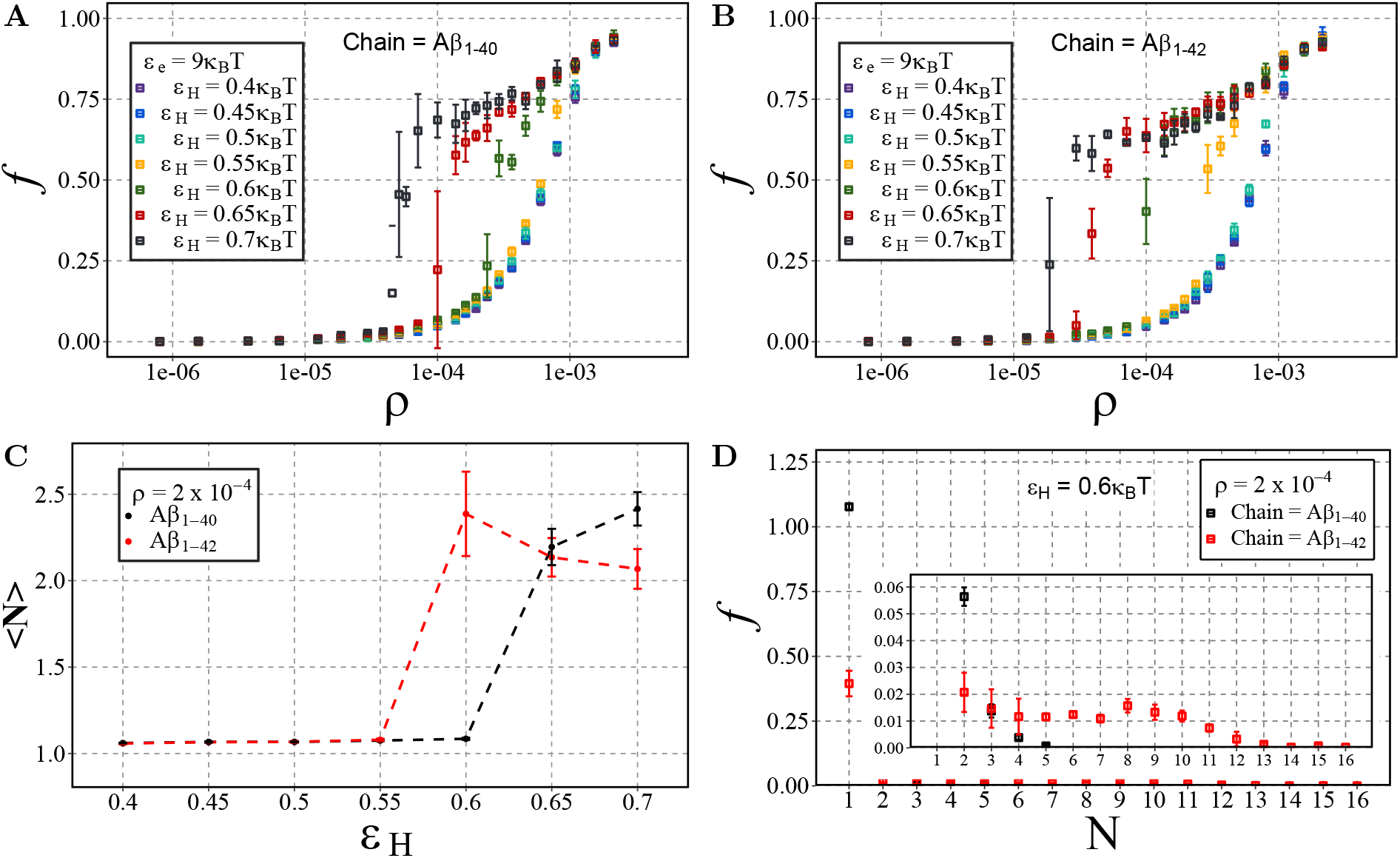
A*β* aggregation with dimerisation (nucleation) energy *ε*_n_ = 3 and elongation energy *ε*_e_ = 9. Each point represents the mean and standard deviation of 3 simulations. The fraction of aggregated material, *f*, is plotted against the number density, *ρ* (in units of the lattice spacing), for varying values of *ε*_H_ using either **(A)** A*β*_1–40_ and **(B)** A*β*_1–42_. **(C)** Displays the change in mean cluster size, 〈*N*〉, against varying hydrophobic interaction energy, *ε*_H_, at a concentration of *ρ* = 2 · 10^−4^. **(D)** A size distribution plot with the fraction of material, *f*, plotted against the cluster size, *N*, with a fixed hydrophobicity of *ε*_H_ = 0.6 at a concentration of *ρ* = 2 · 10^−4^.

### C. Aggregation of Mixed A*β*_1–40_ and A*β*_1–42_

The difference in the critical concentration for condensation of A*β*_1–40_ and A*β*_1–42_, begs the question to what extent our simple model can capture the reported observation of A*β*_1–42_-enhanced aggregation in mixtures of A*β*_1–40_ and A*β*_1–42_ [32, 37]. To investigate this, we have carried out *in silico* mixing experiments for two different values of hydrophobicity. Firstly, we have chosen *ε*_H_ = 0.4*k*_B_*T* as it previously appeared to not have lead to different aggregation dynamics of A*β*_1–42_ and A*β*_1–40_. Secondly, we have chosen *ε*_H_ = 0.6*k*_B_*T*, as this this value showed the largest difference in the aggregation propensity of the two chains, see **Figure 6**.

In **Figure 7** we plot the fraction of aggregated material (panel **A**) and the mean aggregate size (panel **B**) against the fraction of A*β*_1–42_ in a mixture of A*β*_1–42_ and A*β*_1–40_. Having in mind that for strong hydrophobicity structural coarsening renders the system out-of-equilibrium even at long time scales, and thermal equilibrium may never be reached (see **Figure 4**), we have plotted the results after time 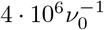 and after 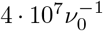. As expected, for low hydrophobicity (*ε*_H_ = 0.4*k*_B_*T*), almost no aggregation takes place and the fraction of aggregated materials is ≈ 0.1, independently of the fraction of long chains. However, for higher hydrophobicity (ε_H_ = 0.6*k*_B_*T*) aggregation occurs rapidly for mixtures with 20% A*β*_1–42_ (solid lines), while at long timescale also mixtures with 10% A*β*_1–42_ start to aggregate (dotted lines). In qualitative agreement with the experimental observations [90], the critical point is strongly biased to a low fraction of A*β*_1–42_ in the mixture (< 10% in our simulations). The slow dynamics is expected, as it is typical for first-order phase separation near the critical point [70]. However, we observe another slow process beyond the critical point, namely the increase of the aggregate size at high concentrations of A*β*_1–42_ (panel **B**). We attribute these to rare events of aggregate fusion.

**FIG. 7:**
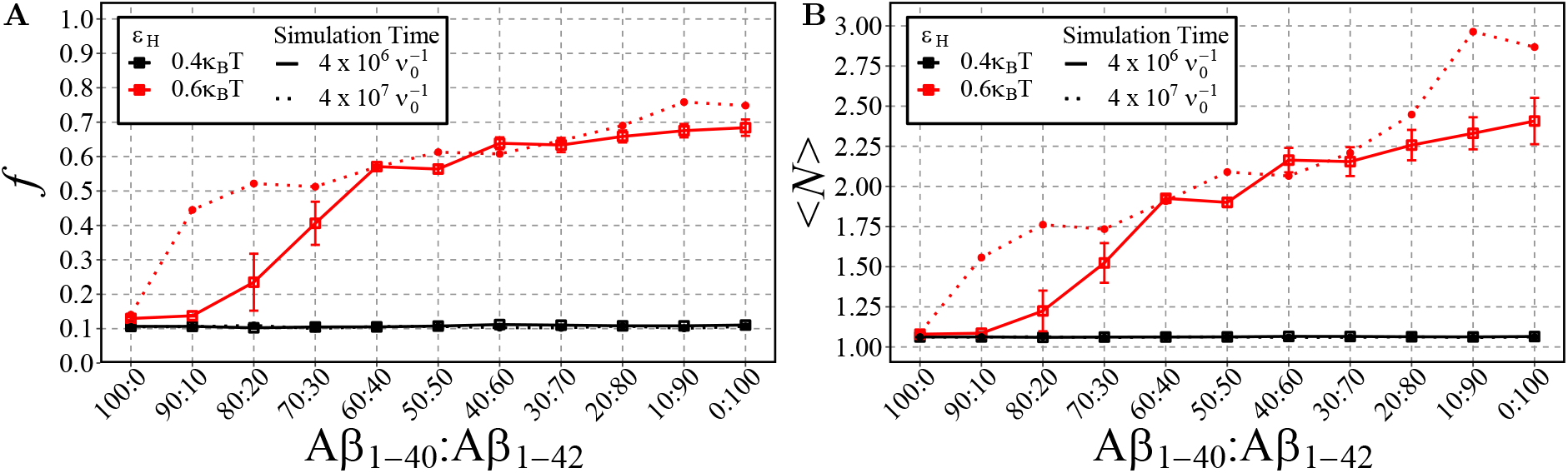
Shown are the fraction of aggregated material, *f*, **(A)** and the mean cluster size, 〈*N*〉, **(B)** using different ratios of A*β*_1–40_ and A*β*_1–42_ chains for two different values of *ε*_H_. 100 total chains were used in each simulation. A consistent box size of 80 × 80 × 80 (approximatly 200uM) was used in all simulations, with *ε*_e_ = 9*k*_B_*T* and *ε*_n_ = 3*k*_B_*T*. For typical simulation times (4 · 10^6^ in units of 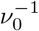; reached in 5 · 10^10^ timesteps), each point represents the mean and standard error of 3 simulations. For longer simulation times 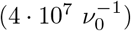, each data point represents the result from a single simulation.

## IV. DISCUSSION AND CONCLUSION

Our study has contributed a molecular modelling approach to address (1) the observation of A*β*_1–42_-enhanced toxic plaques associated with Alzheimer’s disease, despite being the less concentrated species of A*β*, and (2) the hypothesis of condensate-precursors for fibrilation. In our approach we have modelled A*β* as an intrinsically disordered protein with a ‘sticky’ unit that can reversibly dimerise and grow into linear oligomers, and with ‘linker’-type chains of which the beads have either hydrophylic or hydrophobic properties.

The model predicts a rich range of phenomena due to the interplay between the nucleation and elongation of aggregates with the LLPS of condensates, driven by hydrophobic or other non-specific interactions. The high concentration inside the condensates enhances the rate of nucleation through the formation of dimers. Further, at these high concentrations the excluded-volume interactions, which in dilute conditions would hamper the growth of aggregates, are screened, such that the cooperativity of oligomerisation is enhanced. Consequently, we find a sharp transition from the unbound to the aggregated state. However, the dynamics by which aggregates may form is slow close to the critical conditions for LLPS. At concentrations above the critical concentration, the growth of aggregates is slow due to the relatively slow dynamics of Ostwald ripening and/or fusion events.

A similar secondary-nucleation mechanism was discussed previously [91, 92], and it was found that the rate constants for primary nucleation, elongation and secondary nucleation are 100-fold, 10-fold and 3-fold greater respectively for A*β*_1–42_ compared to A*β*_1–40_ [93]. This provides evidence that the addition of two c-terminal hydrophobic amino acids promotes aggregation propensity, in particular the rate of primary nucleation. This is in agreement with [94] who propose that primary nucleation of A*β* is driven by non-specific hydrophobic interactions which explains the difference in the aggregation rates of A*β*_1–40_ and A*β*_1–42_.

Despite its simplicity, the model also predicts that a small fraction (< 10%) of A*β*_1–42_ in a mixture of A*β*_1–42_ and A*β*_1–40_ shifts the critical concentration for the LLPS of condensates to lower values, which in turn leads to the nucleated self-assembly of fibrillar oligomers at reduced concentrations. This finding is qualitatively consistent with [90] who found that even a small change from a 1:9 to a 3:7 ratio of A*β*_1–42_:A*β*_1–40_ caused a dramatic change in the aggregation kinetics and toxicity of the two mixtures *in vitro* and *in vivo.* This 3:7 ratio is of particular importance as it reflects the ratio of A*β*_1–42_: A*β*_1–40_ observed in familial AD patients [30] suggesting that maintaining a physiological ratio of A*β*_1–42_: A*β*_1–40_ is of great importance and may be an effective therapeutic target.

Whether A*β*_1–40_ and A*β* _1-42_ can co-fibrilise is still under debate, [95] found that mixing A*β*_1–40_ and A*β*_1–42_ leads to the generation of separate homomolecular fibrils. In contrast, studies have showed that mixing A*β*_1–40_ and A*β*_1–42_ leads to the formation of mixed oligomers and fibrils [29, 96]. However, it is accepted that A*β*_1–40_ and A*β*_1–42_ interact at the molecular level with increased levels of A*β*_1–40_ inhibiting fibril formation and increased levels of A*β*_1–42_ promoting aggregation [97–100]. Despite studies disagreeing on co-fibrilisation, they agree that prefibrillar intermediates consisting of both A*β*_1–40_ and A*β*_1–42_ exist. Our findings suggest that A*β*_1–42_ interacts with A*β*_1–40_, facilitating its aggregation. When only A*β*_1–40_ chains are present and *ε*_H_ = 0.6*k*_B_*T*, approximately 10% of the chains are aggregated. However, with 4 · 10^6^ 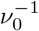, when A*β*_1–40_ and A*β*_1–42_ are mixed at a 60:40 ratio, f increases to approximately 60%. If only A*β*_1–42_ chains are capable of aggregation, *f* = 40% at maximum. As this is not the case, it demonstrates that addition of A*β* _1-42_ is sufficient to promote the aggregation of A*β*_1–40_ under these conditions. One explanation may be that when the more aggregation prone A*β*_1–42_ is present, it forms the primary nuclei overcoming the initial energy barrier that then enables A*β*1-40 to aggregate. This theory is supported by [95] who showed that upon mixing A*β*_1–42_ and A*β*_1–40_ it is A*β*_1–42_ that aggregates first, followed by A*β*1-40.

In conclusion, we have presented a simple sticker-and-linker lattice model that captures a wide range of molecular phenomena of relevance to the literature on A*β* aggregation, and which enables us to interpret those phenomena in terms of the simple concepts of nucleation-elongation models and of hydrophobicity and polymeric excluded-volume effects at the level of coarse-grained sticky-polymer models. We hope this approach will lead to further (quantitative) refinements to the understanding of typical experimental time- and length scales of A*β* aggregation in few-component *in vitro* and complex multi-component *in vivo* studies.

## ACKNOWLEDGMENTS

JC acknowledges financial support by the Physics of Life UK. SQ acknowledges support from Alzheimer’s Research UK (RF2019-A-001). CS acknowledges support from the Engineering and Physical Sciences Research Council [grant number EPSRC (EP/N031431/1)]. CS thanks Dr. Ed Higgins, Mitch Aitchison, Thibault Delpech and Arsenii Zats for their assistance in the development and implementation of the model. CS and JC are grateful for computational support from the University of York Research and High Performance Computing service, The Viking Cluster.

## Notes

### Competing Interest Statement

The authors have declared no competing interest.

## References

[1] A. A. D. T. Abeysinghe, R. D. U. S. Deshapriya, and C. Udawatte, Life Sciences 256, 117996 (2020).

[2] Q. Zhang, J. Sidorenko, B. Couvy-Duchesne, R. E. Marioni, M. J. Wright, A. M. Goate, E. Marcora, K.-l. Huang, T. Porter, S. M. Laws, P. S. Sachdev, K. A. Mather, N. J. Armstrong, A. Thalamuthu, H. Brodaty, L. Yengo, J. Yang, N. R. Wray, A. F. McRae, and P. M. Visscher, Nature Communications 11, 4799 (2020), number: 1 Publisher: Nature Publishing Group.

[3] N. Zhao, Y. Ren, Y. Yamazaki, W. Qiao, F. Li, L. M. Felton, S. Mahmoudiandehkordi, A. Kueider-Paisley, B. Sonoustoun, M. Arnold, F. Shue, J. Zheng, O. N. Attrebi, Y. A. Martens, Z. Li, L. Bastea, A. D. Meneses, K. Chen, J. W. Thompson, L. St John-Williams, M. Tachibana, T. Aikawa, H. Oue, L. Job, A. Yamazaki, C.-C. Liu, P. Storz, Y. W. Asmann, N. Ertekin-Taner, T. Kanekiyo, R. Kaddurah-Daouk, and G. Bu, Neuron 106, 727 (2020).

[4] T. Ayodele, E. Rogaeva, J. T. Kurup, G. Beecham, and C. Reitz, Current Neurology and Neuroscience Reports 21, 4 (2021).

[5] C. Patterson, (2018).

[6] R. A. Stelzmann, H. Norman Schnitzlein, and F. Reed Murtagh, Clinical Anatomy 8, 429 (1995), _eprint: https://onlinelibrary.wiley.com/doi/pdf/10.1002/ca.980080612.

[7] Z. Breijyeh and R. Karaman, Molecules 25, 5789 (2020), number: 24 Publisher: Multidisciplinary Digital Publishing Institute.

[8] G.-f. Chen, T.-h. Xu, Y. Yan, Y.-r. Zhou, Y. Jiang, K. Melcher, and H. E. Xu, Acta Pharmacologica Sinica 38, 1205 (2017).

[9] T. Guo, D. Zhang, Y. Zeng, T. Y. Huang, H. Xu, and Y. Zhao, Molecular Neurodegeneration 15, 40 (2020).

[10] H. Hampel, J. Hardy, K. Blennow, C. Chen, G. Perry, S. H. Kim, V. L. Villemagne, P. Aisen, M. Vendruscolo, T. Iwatsubo, C. L. Masters, M. Cho, L. Lannfelt, J. L. Cummings, and A. Vergallo, Molecular Psychiatry 26, 5481 (2021), number: 10 Publisher: Nature Publishing Group.

[11] Y.-w. Zhang, R. Thompson, H. Zhang, and H. Xu, Molecular Brain 4, 3 (2011).

[12] C. Haass, C. Kaether, G. Thinakaran, and S. Sisodia, Cold Spring Harbor Perspectives in Medicine 2, a006270 (2012), publisher: Cold Spring Harbor Laboratory Press.

[13] J. A. Hardy and G. A. Higgins, Science 256, 184 (1992), publisher: American Association for the Advancement of Science.

[14] E. Karran, M. Mercken, and B. D. Strooper, Nature Reviews Drug Discovery 10, 698 (2011), number: 9 Publisher: Nature Publishing Group.

[15] R. J. Castellani, G. Plascencia-Villa, and G. Perry, Laboratory Investigation 99, 958 (2019), number: 7 Publisher: Nature Publishing Group.

[16] F. Kametani and M. Hasegawa, Frontiers in Neuroscience 12 (2018).

[17] E. M. Arenaza-Urquijo and P. Vemuri, Neurology 90, 695 (2018), publisher: Wolters Kluwer Health, Inc. on behalf of the American Academy of Neurology Section: Views & amp; Reviews.

[18] B. Dubois, N. Villain, G. B. Frisoni, G. D. Rabinovici, M. Sabbagh, S. Cappa, A. Bejanin, S. Bombois, S. Epelbaum, M. Teichmann, M.-O. Habert, A. Nordberg, K. Blennow, D. Galasko, Y. Stern, C. C. Rowe, S. Salloway, L. S. Schneider, J. L. Cummings, and H. H. Feldman, The Lancet Neurology 20, 484 (2021).

[19] R. Ricciarelli and E. Fedele, Current Neuropharmacology 15, 926 (2017).

[20] M. Tolar, S. Abushakra, and M. Sabbagh, Alzheimer’s & Dementia (2019), 10.1016/j.jalz.2019.09.075.

[21] S. M. Chafekar, F. Baas, and W. Scheper, Biochimica et Biophysica Acta (BBA) - Molecular Basis of Disease 1782, 523 (2008).

[22] K. Ono, M. M. Condron, and D. B. Teplow, Proceedings of the National Academy of Sciences 106, 14745 (2009), publisher: Proceedings of the National Academy of Sciences.

[23] U. Sengupta, A. N. Nilson, and R. Kayed, EBioMedicine 6, 42 (2016).

[24] Y.-r. Huang and R.-t. Liu, International Journal of Molecular Sciences 21, 4477 (2020), number: 12 Publisher: Multidisciplinary Digital Publishing Institute.

[25] D. J. Selkoe and J. Hardy, EMBO molecular medicine 8, 595 (2016).

[26] S. A. Ghadami, S. Chia, F. S. Ruggeri, G. Meisl, F. Bemporad, J. Habchi, R. Cascella, C. M. Dobson, M. Vendruscolo, T. P. J. Knowles, and F. Chiti, Biomacromolecules 21, 1112 (2020), publisher: American Chemical Society.

[27] A. Tamaoka, T. Kondo, A. Odaka, N. Sahara, N. Sawamura, K. Ozawa, N. Suzuki, S. Shoji, and H. Mori, Biochemical and Biophysical Research Communications 205, 834 (1994).

[28] T. Iwatsubo, A. Odaka, N. Suzuki, H. Mizusawa, N. Nukina, and Y. Ihara, Neuron 13, 45 (1994).

[29] L. Gu and Z. Guo, Biochemical and Biophysical Research Communications 534, 292 (2021).

[30] D. Scheuner, C. Eckman, M. Jensen, X. Song, M. Citron, N. Suzuki, T. D. Bird, J. Hardy, M. Hutton, W. Kukull, E. Larson, E. Levy-Lahad, M. Viitanen, E. Peskind, P. Poorkaj, G. Schellenberg, R. Tanzi, W. Wasco, L. Lannfelt, D. Selkoe, and S. Younkin, Nature Medicine 2, 864 (1996).

[31] S. Hecimovic, J. Wang, G. Dolios, M. Martinez, R. Wang, and A. M. Goate, Neurobiology of Disease 17, 205 (2004).

[32] S. Kumar-Singh, J. Theuns, B. Van Broeck, D. Pirici, K. Vennekens, E. Corsmit, M. Cruts, B. Dermaut, R. Wang, and C. Van Broeckhoven, Human Mutation 27, 686 (2006), _eprint: https://onlinelibrary.wiley.com/doi/pdf/10.1002/humu.20336.

[33] L. Chávez-Gutiérrez and M. Szaruga, Seminars in Cell & Developmental Biology Gamma Secretase, 105, 75 (2020).

[34] I. Baldeiras, I. Santana, M. J. Leitão, H. Gens, R. Pascoal, M. Tábuas-Pereira, J. Beato-Coelho, D. Duro, M. R. Almeida, and C. R. Oliveira, Alzheimer’s Research & Therapy 10, 33 (2018).

[35] S. Lehmann, C. Delaby, G. Boursier, C. Catteau, N. Ginestet, L. Tiers, A. Maceski, S. Navucet, C. Paquet, J. Dumurgier, E. Vanmechelen, H. Vanderstichele, and A. Gabelle, Frontiers in Aging Neuroscience 10 (2018).

[36] A. Nakamura, N. Kaneko, V. L. Villemagne, T. Kato, J. Doecke, V. Doré, C. Fowler, Q.-X. Li, R. Martins, C. Rowe, T. Tomita, K. Matsuzaki, K. Ishii, K. Ishii, Y. Arahata, S. Iwamoto, K. Ito, K. Tanaka, C. L. Masters, and K. Yanagisawa, Nature 554, 249 (2018), number: 7691 Publisher: Nature Publishing Group.

[37] H. Zetterberg, Journal of Neuroscience Methods Methods and Models in Alzheimer’s Disease Research, 319, 2 (2019).

[38] J. D. Doecke, V. Pérez-Grijalba, N. Fandos, C. Fowler, V. L. Villemagne, C. L. Masters, P. Pesini, and M. Sarasa, Neurology 94, e1580 (2020).

[39] Y. A. R. Mahaman, K. S. Embaye, F. Huang, L. Li, F. Zhu, J.-Z. Wang, R. Liu, J. Feng, and X. Wang, Ageing Research Reviews 74, 101544 (2022).

[40] C. E. Teunissen, I. M. W. Verberk, E. H. Thijssen, L. Vermunt, O. Hansson, H. Zetterberg, W. M. van der Flier, M. M. Mielke, and M. del Campo, The Lancet Neurology 21, 66 (2022).

[41] N. Fandos, V. Pérez-Grijalba, P. Pesini, S. Olmos, M. Bossa, V. L. Villemagne, J. Doecke, C. Fowler, C. L. Masters, and M. Sarasa, Alzheimer’s & Dementia: Diagnosis, Assessment & Disease Monitoring 8, 179 (2017).

[42] K. Yaffe, A. Weston, N. R. Graff-Radford, S. Satterfield, E. M. Simonsick, S. G. Younkin, L. H. Younkin, L. Kuller, H. N. Ayonayon, J. Ding, and T. B. Harris, JAMA :the journal of the American Medical Association 305, 261 (2011).

[43] V. Pérez-Grijalba, J. Romero, P. Pesini, L. Sarasa, I. Monleón, I. San-José, J. Arbizu, P. Martínez-Lage, J. Munuera, A. Ruiz, L. Tárraga, M. Boada, M. Sarasa, and The AB255 Study Group, The Journal of Prevention of Alzheimer’s Disease 6, 34 (2019).

[44] Y. Y. Lim, P. Maruff, N. Kaneko, J. Doecke, C. Fowler, V. L. Villemagne, T. Kato, C. C. Rowe, Y. Arahata, S. Iwamoto, K. Ito, K. Tanaka, K. Yanagisawa, C. L. Masters, and A. Nakamura, Journal of Alzheimer’s Disease 77, 1057 (2020), publisher: IOS Press.

[45] K. V. Giudici, P. de Souto Barreto, S. Guyonnet, Y. Li, R. J. Bateman, B. Vellas, and MAPT/DSA Group, JAMA Network Open 3, e2028634 (2020).

[46] Y. Shin and C. P. Brangwynne, Cellular Biophysics 357, 1253 (2017).

[47] X. Jin, J.-E. Li, C. Schaefer, X. Luo, X. Luo, A. J. M. Wollman, T. Tian, X. Zhang, X. Chen, Y. Li, Y. Pu, T. C. B. McLeish, M. C. Leake, and F. Bai, Sci. Adv. 7, eabh2929 (2021).

[48] S. Wegmann, B. Eftekharzadeh, K. Tepper, K. M. Zoltowska, R. E. Bennett, S. Dujardin, P. R. Laskowski, D. MacKenzie, T. Kamath, C. Commins, C. Vanderburg, A. D. Roe, Z. Fan, A. M. Molliex, A. Hernandez-Vega, D. Muller, A. A. Hyman, E. Mandelkow, J. P. Taylor, and B. T. Hyman, The EMBO Journal 37, e98049 (2018), https://www.embopress.org/doi/pdf/10.15252/embj.201798049.

[49] L. Leibler, M. Rubinstein, and R. H. Colby, Macromolecules 24, 4701 (1991).

[50] C. Schaefer, P. R. Laity, C. Holland, and T. C. B. McLeish, Macromolecules 53, 2669 (2020).

[51] C. Schaefer and T. C. B. McLeish, J. Rheol. 66, 515 (2022).

[52] J.-M. Choi, A. S. Holehouse, and R. V. Pappu, Annu. Rev. Biophys. 49, 107 (2020).

[53] J. R. Gissinger, B. D. Jensen, and K. E. Wise, Polymer 128, 211 (2017).

[54] G. Cui, V. A. H. Boudara, Q. Huang, G. P. Baeza, A. J. Wilson, O. Hassager, D. J. Read, and J. Mattsson, J. Rheol. 62, 1155 (2018).

[55] C. Raffaelli, A. Bose, C. H. M. P. Vrusch, S. Ciarella, T. Davris, N. B. Tito, A. V. Lyulin, W. G. Ellenbroek,, and C. Storm, in Self-Healing and Self-Recovering Hydrogels, Vol. 285, edited by C. Creton and O. Okay (CRC Press, 2020).

[56] I. Carmesin and K. Kremer, Macromolecules 21, 273 (1988).

[57] I. Carmesin and K. Kremer, J. de Physique 51, 915 (1990).

[58] M. A. Bates, J. Chem. Phys 120, 3986 (2002).

[59] M. Feric, N. Vaidya, T. S. Harmon, D. M. Mitrea, L. Zhu, T. M. Richardson, R. W. Kriwacki, R. V. Pappu, and C. P. Brangwynne, Cell 165, 1686 (2016).

[60] T. S. Harmon, A. S. Holehouse, M. K. Rosen, and R. V. Pappu, eLife 6, e30294 (2017).

[61] E. Reister, M. Müller, and K. Binder, Phys. Rev E 64, 041804 (2001).

[62] E. Reister and M. Müller, J. Chem. Phys. 118, 8476 (2003).

[63] G. Subramanian and S. Shanbhag, Phys. Rev. E 80, 041806 (2009).

[64] H. L. Trautenberg, J.-U. Sommer, and D. Goritz, J. Chem. Soc. Faraday Trans. 91, 2649 (1995).

[65] J. J. Lukkien, J. P. L. Segers, P. A. J. Hilbers, R. J. Gelten, and A. P. J. Jansen, Physical Review E 58, 2598 (1998).

[66] J. Danielsson, J. Jarvet, P. Damberg, and A. Gräslund, The FEBS Journal 272, 3938 (2005).

[67] J. Roche, Y. Shen, J. H. Lee, J. Ying, and A. Bax, Biochemistry 55, 762 (2016), publisher: American Chemical Society.

[68] M. A. Wälti, J. Orts, B. Vögeli, S. Campioni, and R. Riek, ChemBioChem 16, 659 (2015), _eprint: https://onlinelibrary.wiley.com/doi/pdf/10.1002/cbic.201402595.

[69] X. Zhu, R. P. Bora, A. Barman, R. Singh, and R. Prabhakar, The Journal of Physical Chemistry B 116, 4405 (2012).

[70] P. C. Hohenberg and B. I. Halperin, Rev. of Mod. Phys. 49, 435 (1977).

[71] C. Schaefer, Phys. Rev. Lett. 120, 036001 (2018).

[72] C. Schaefer, S. Paquay, and T. C. B. McLeish, Soft Matter 15, 8450 (2019).

[73] K. Binder, Colloid & Polymer Sci. 265, 273 (1987).

[74] M. Saito and M. Matsumoto, in Monte Carlo and Quasi-Monte Carlo Methods 2006, edited by A. Keller, S. Heinrich, and H. Niederreiter (Springer Berlin Heidelberg, Berlin, Heidelberg, 2008) pp. 607–622.

[75] S. Müller-Späth, A. Soranno, V. Hirschfeld, H. Hofmann, S. Rüegger, L. Reymond, D. Nettels, and B. Schuler, PNAS 107, 14609 (2010).

[76] R. Wuttke, H. Hofmann, D. Nettels, M. B. Borgia, J. Mittal, R. B. Best, and B. Schuler, PNAS 11, 5213 (2014).

[77] W. Paul, M. Muller, K. Binder, M. R. Stukan, and V. A. Ivanov, P. Pasini et al. (eds.), Computer Simulations of Liquid Crystals and Polymers: Monte Carlo simulations of semi-flexible polymers (Kluwer Academic Publisher, NL, 2005) pp. 171–190.

[78] M. O. Steinhauser, J. Chem. Phys. 122, 094901 (2005).

[79] P. Hortschansky, V. Schroeckh, T. Christopeit, G. Zandomeneghi, and M. Fändrich, Protein Sci. 14, 1753 (2005).

[80] M. Novo, S. Freire, and W. Al-Soufi, Sci. Reports 8, 1783 (2018).

[81] A. J. Bray, Adv. Phys. 51, 481 (2002).

[82] A. Singh, S. Puri, and C. Dasgupta, J. Phys. Chem. B 116, 4519 (2012).

[83] R. B. Martin, Chem. Rev. 96, 3044 (1996).

[84] P. van der Schoot, in Supramolecular polymers (2nd Edition), edited by A. Ciferri (CRC Press, 2005) pp. 77–106.

[85] T. F. A. de Greef, M. M. J. Smulders, M. Wolffs, A. P. H. J. Schenning, R. P. Sijbesma, and E. W. Meijer, Chem. Rev. 109, 5687 (2009).

[86] M. E. Cates and S. J. Candau, J. Phys.: Condens. Matter 2, 6869 (1990).

[87] C. Kulkarni, E. W. Meijer, and A. R. A. Palmans, Acc. Chem. Res. 50, 1928 (2017).

[88] H. Weber, W. Paul, and K. Binder, Phys. Rev. E 59, 2168 (1999).

[89] M. Smulders, M. Nieuwenhuizen, T. de Greef, P. van der Schoot, A. Schenning, and E. Meijer, Chemistry - A European Journal 16, 362 (2010), https://chemistry-europe.onlinelibrary.wiley.com/doi/pdf/10.1002/chem.200902415.

[90] I. Kuperstein, K. Broersen, I. Benilova, J. Rozenski, W. Jonckheere, M. Debulpaep, A. Vandersteen, I. Segers-Nolten, K. Van Der Werf, V. Subramaniam, D. Braeken, G. Callewaert, C. Bartic, R. D’Hooge, I. C. Martins, F. Rousseau, J. Schymkowitz, and B. De Strooper, The EMBO journal 29, 3408 (2010).

[91] T. C. T. Michaels, A. Saric, S. Curk, K. Bernfur, P. Arosio, G. Meisl, A. J. Dear, S. I. A. Cohen, C. M. Dobson, M. Vendruscolo, S. Linse, and T. P. J. Knowles, Nature Chemistry 12, 445 (2020), number: 5 Publisher: Nature Publishing Group.

[92] S. I. A. Cohen, S. Linse, L. M. Luheshi, E. Hellstrand, D. A. White, L. Rajah, D. E. Otzen, M. Vendruscolo, C. M. Dobson, and T. P. J. Knowles, Proceedings of the National Academy of Sciences of the United States of America 110, 9758 (2013).

[93] G. Meisl, X. Yang, E. Hellstrand, B. Frohm, J. B. Kirkegaard, S. I. A. Cohen, C. M. Dobson, S. Linse, and T. P. J. Knowles, Proceedings of the National Academy of Sciences 111, 9384 (2014), publisher: Proceedings of the National Academy of Sciences.

[94] J. Roche, Y. Shen, J. H. Lee, J. Ying, and A. Bax, Biochemistry 55, 762 (2016), publisher: American Chemical Society.

[95] R. Cukalevski, X. Yang, G. Meisl, U. Weininger, K. Bernfur, B. Frohm, T. P. J. Knowles, and S. Linse, Chemical Science 6, 4215 (2015), publisher: The Royal Society of Chemistry.

[96] L. Cerofolini, E. Ravera, S. Bologna, T. Wiglenda, A. Böddrich, B. Purfüst, I. Benilova, M. Korsak, G. Gallo, D. Rizzo, L. Gonnelli, M. Fragai, B. D. Strooper, E. E. Wanker, and C. Luchinat, Chemical Communications 56, 8830 (2020), publisher: The Royal Society of Chemistry.

[97] K. Hasegawa, I. Yamaguchi, S. Omata, F. Gejyo, and H. Naiki, Biochemistry 38, 15514 (1999).

[98] Y. Yan and C. Wang, Journal of Molecular Biology 369, 909 (2007).

[99] A. Jan, O. Gokce, R. Luthi-Carter, and H. A. Lashuel, Journal of Biological Chemistry 283, 28176 (2008), publisher: Elsevier.

[100] K. Pauwels, T. L. Williams, K. L. Morris, W. Jonckheere, A. Vandersteen, G. Kelly, J. Schymkowitz, F. Rousseau, A. Pastore, L. C. Serpell, and K. Broersen, The Journal of Biological Chemistry 287, 5650 (2012).

